# Prophage maintenance is determined by environment-dependent selective sweeps rather than mutational availability

**DOI:** 10.1101/2023.03.21.533645

**Authors:** Zachary M. Bailey, Claudia Igler, Carolin C. Wendling

**Author notes:** **Corresponding authors:** Zachary Bailey, Carolin Wendling. **Author contributions:** CCW and CI obtained funding. CCW and ZB designed the experiment. ZB performed the experiment and analyzed the experimental and genomic data. ZB and CI developed the model. ZB ran the simulations. All authors wrote the manuscript and approved the final version of this manuscript. **Acknowledgements:** We thank the members of the Pathogen Ecology group at ETH Zurich for their productive discussions especially Ricardo Leon-Sampedro, Ayush Pathak and Mathilde Boumasmoud. We thank Rahel Bruhlman for assistance in performing PCRs, and Christopher Witzany and Daniel Angst for lending cluster computational time. **Data accessibility statement:** The underlying data for this study is available at https://datadryad.org/stash/share/5nF31yo9SkBCpyQ6ZYU8OF-s-JhQluCxRUUSEuTLcGY. Genomic information for this study is available under study accession number PRJEB60436 at the European Nucleotide Archive.

## Abstract

Prophages, viral sequences integrated into bacterial genomes, can be beneficial and costly. Despite the risk of prophage activation and subsequent bacterial death, active prophages are present in most bacterial genomes. However, our understanding of the selective forces that maintain prophages in bacterial populations is limited. Combining experimental evolution with stochastic modelling, we show that prophage maintenance and loss are primarily determined by environmental conditions that alter the net fitness effect of a prophage. When prophages are too costly, they are rapidly lost through environment-specific sequences of selective sweeps. Conflicting selection pressures that select against the prophage but for a prophage-encoded accessory gene can maintain prophages. The dynamics of prophage maintenance additionally depends on the sociality of this accessory gene. Non-cooperative genes maintain prophages at higher frequencies than cooperative genes, which can protect phage-free ‘cheaters’ that may emerge if prophage costs outweigh their benefits. Our simulations suggest that environmental variation plays a larger role than mutation rates in determining prophage maintenance. These findings highlight the complexity of selection pressures that act on mobile genetic elements and challenge our understanding of the role of environmental factors relative to random chance events in shaping the evolutionary trajectory of bacterial populations.

## Introduction

Prophages, viruses integrated into bacterial genomes, pose a challenging conflict for their hosts. The ability of a prophage to revert to the lytic cycle - which leads to the production of new phage particles subsequently lysing and killing the bacterial cell, puts their hosts at constant risk of death. Yet, their presence in bacterial genomes can increase bacterial fitness in several ways. Firstly, lysogens, i.e., cells carrying one or more prophages, are protected from superinfection of bacteriophages with the same immunity class through superinfection exclusion provided by the integrated prophage (1). Secondly, prophages can encode accessory genes, which can change their host’s phenotype through a process known as lysogenic conversion (2). The expression of these genes, whose products often confer a cooperative benefit (3), can increase the fitness of lysogens and potentially surrounding bacteria in certain environments. Lastly, prophages can switch back to the lytic cycle, during which they replicate within the bacterial host cell, the subsequently released phage particles can kill phage-susceptible competitors which enhances the fitness of the remaining lysogen population (4,5).

However, the metabolic burden of viral protein expression during lysogenic conversion can render the respective prophage costly in certain environments (6,7). Furthermore, the induction of the lytic cycle, which can occur spontaneously or in response to external DNA-damaging stressors (8–11), inevitably results in the death of the bacterial cell. Depending on the rate of induction, i.e., the fraction of the bacterial population that undergoes lysis, such events can drive entire lysogen populations to extinction (personal observation, Supp. Fig. S1).

Bacterial populations can mitigate costs imposed by prophages in several ways, including the accumulation of deleterious mutations in the prophage (12–14) or mutations in the bacterial genome that alter phage induction rate (15). Alternatively, bacteria can eliminate prophages from their genomes (12,13,16,17) following incomplete activation of the prophage or through partial or complete deletions (18,19).

Despite these mechanism to ameliorate the costs of prophages, the persistence of functional prophages in a majority of bacterial genomes - up to 83% (20) – raises important evolutionary questions about the selective forces that maintain or abolish prophages in nature. Specifically, it remains unclear under which conditions prophages become costly enough to be lost from a bacterial population, and under which conditions they provide enough benefits to be maintained. We predict that prophage maintenance and the evolutionary trajectory of lysogen populations are influenced by the frequency of prophage induction and any additional costs and benefits associated with the prophage, all of which are subject to variation across different environments (7).

To explore the dynamics of prophage maintenance, we co-evolved lysogenic *E. coli* populations carrying Lambda prophages, under selection pressures that altered the net fitness effect of prophage carriage. We found that high rates of prophage induction selected for fast prophage loss driven by strong directional selection, resulting in environment-dependent selective sweeps. The first was a transient sweep during which bacterial populations acquired chromosomal phage resistance to prevent reinfections by free phages. The second sweep was characterized by an increasing frequency of phage-free cells, with a phage-resistant and phage-free genotype rapidly reaching fixation after approximately 500 generations. Combining these data with an empirically parameterized model, we found that conflicting selection pressures and non-cooperative benefits provided by the prophage resulted in slower and more stochastic prophage loss, even when prophages were costly. Using linear discriminant analysis, we found that prophage loss is largely driven by environmental factors such as antibiotic concentration or prophage induction, rather than mutation rates. This implies that prophage loss is not simply a matter of random mutations but is influenced by a complex interplay between environmental factors and environment-specific genetic changes. Given the omnipresence of prophages in the microbial world, our findings suggest that prophages play an integral role in microbial evolution because their maintenance is driven by environmental variation and genetically determined social interactions.

## Methods

### 1. Study System

We used *Escherichia coli* K-12 MG1655, which is cured of phage Lambda and contains a constitutively expressed YFP marker to construct two different lysogens. One lysogen contained Lambda+ which encodes a constitutively expressed Ampicillin (Amp) resistance gene (BlaTEM116) (7). The other lysogen contained the same prophage yet lacked the resistance gene. Lysogenization was confirmed by PCR using primers that target the attB site in *E. coli* and the *int* gene of Lambda+ (21). Hereafter, we refer to lambda free bacteria as non-lysogens (*n_lys)*, to lysogenized bacteria without the resistance gene as susceptible lysogens (*s_lys)*, and to lysogenized bacteria with the Amp-resistance gene (Amp^R^) as resistant lysogens, (*r_lys)*.

### 2. Selection Experiment

#### Experimental design

We initiated a selection experiment to observe prophage maintenance over time in four different selection regimes using three bacterial genotypes, i.e., *n_lys, s_lys* and *r_lys* in a full-factorial design. With six biological replicates, each originating from single colonies, our design resulted in a total of 72 populations. We used the following selection regimes: (i) Amp at IC_50_ (i.e., 6.4 ug/ml) selecting for the prophage encoded Amp^R^ present in the *r_lys* population, (ii) Mitomycin C (MMC) at a concentration of 0.1 ug/ml which induces the lytic cycle of prophages causing massive cell death due to host lysis (Supp. Fig. 1) (22,23) and increases the costs of prophage carriage for the *s_lys* and *r_lys* populations, (iii) a combination of Amp and MMC at the same concentrations as above, to create opposing selective pressures on the *r_lys* populations and two competing selective pressures on *s_lys* populations, and (iv) LB broth as control. Unless otherwise stated, all bacteria were grown at 37° C and constant shaking at 180 rpm.

**Figure 1:**
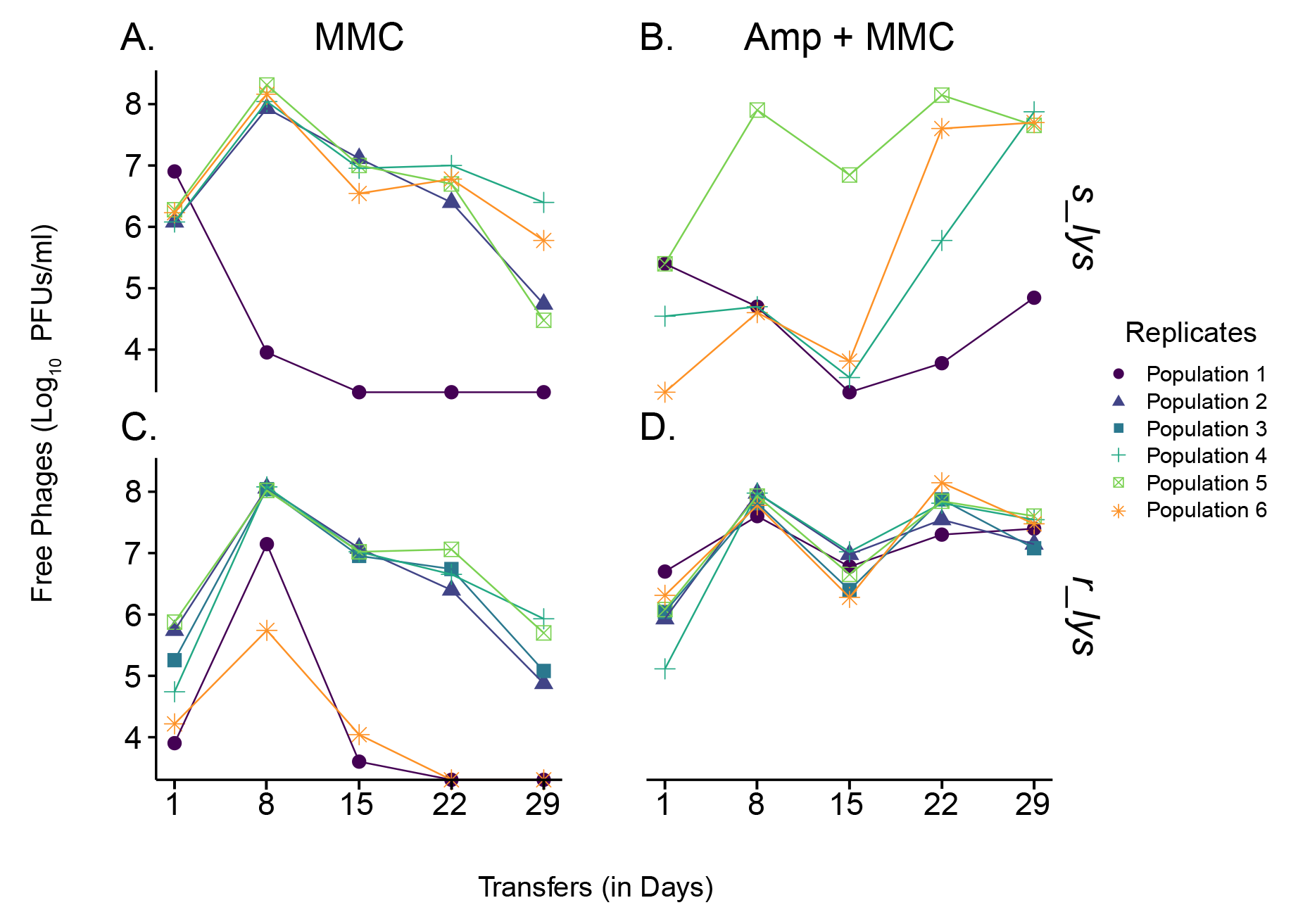
Ecological dynamics of free phage particles in lysogen populations evolved in the presence of MMC (left) or MMC+Amp (right). Log values of the number of phages (Plaque Forming Units, PFUs/ml) are shown for 29 transfers of selection. Phages either carried no Amp-resistance gene (Amp^R^), i.e., susceptible lysogens *s_lys* (top), or carried the corresponding Amp^R^, i.e., resistant lysogens *r_lys* (bottom). Replicate populations are differentiated by symbol and color.

#### Experimental setup

We used 10 µl of an overnight culture to inoculate each treatment. All 72 populations were grown in 96 deep-well plates at a total volume of 1 ml. Every day, we transferred 1% of each population to fresh medium for a total of 30 transfers. After the first 24 hours, and subsequently every 7 transfers (T), we quantified bacterial and phage population dynamics. Additionally, we froze 100 µl of each population every transfer in 25% glycerol at −80° C. Concurrently, every seven transfers, we made glycerol cryo stocks of 24 individual clones from each population which we used for follow-up analyses. At the end of the experiment, we detected contamination in two *s_lys* populations and excluded them from later analysis.

#### Bacterial and phage population dynamics

*Bacterial dynamics:* We measured bacterial cell counts as cells/ml using a Novocyte Novosampler Pro using 1:1000 dilutions of our cell cultures, following a previously established protocol in Wendling et al., 2020 (7).

*Phage dynamics:* We quantified the number of free phages as plaque forming units (PFU/ml) from each population using a plaque spot assay on a lawn of phage susceptible ancestral *E. coli* K-12 MG1655 as described in Wendling et al., 2020 (7). In populations exposed to MMC, we observed a decrease in free phage particles over time (Figure 1). This suggested either (i) a gradual reduction at the population level, resulting from lower burst size and phage productivity, or (ii) the presence of two different phenotypes, of which one stopped producing phages completely while the other produced phages at a rate comparable to the ancestral lysogen. To infer the underlying mechanism, we quantified phage production as PFU/ml, as described above, on ancestral clones and transfers 8, and 29 of 24 randomly selected clones from two *s_lys* and *r_lys* populations originating from either the MMC, or the Amp+MMC treatment. This assay confirmed that the observed decline in free phage particles observed at the population level results from individual clones that did not produce free phages. This suggested that these clones either lost or inactivated their prophages.

### 3. Whole Genome SequencingS

#### Sequencing

To infer the underlying molecular mechanism(s) behind the observed loss in phage particle production of individual clones that evolved in the presence of MMC, we sequenced both single clones and whole populations from the selection experiment.

##### Single clones

We sequenced single clones from the *s_lys* and *r_lys* populations that originated either from (i) LB (ii) the MMC, or the (iii) Amp+MMC treatment. We randomly selected two clones from two replicate populations from T8 and T29 as well as one ancestral clone per genotype.

##### Whole populations

We sequenced whole population samples from all 72 populations from the end of the experiment. Additionally, we included two populations from the Amp+MMC treatment from intermediate time-points, i.e., T8,T15, and T22 to track the presence of candidate mutations across time.

We extracted whole genome DNA using the GenElute Bacterial Genomic DNA extraction kit from Sigma-Aldrich. Library preparation and whole genome sequencing was done at the Functional Genomics Center Zurich (FGCZ) using a NovaSeq Illumina sequencer giving us FASTQ files with an average coverage of over 100 ×.

#### Sequence analysis

##### Individual Clones

We performed a quality check and trimmed Illumina adapter sequences using Trim Galore with FastQC on the trimmed reads (24–26). We analyzed these trimmed and filtered reads using breseq (v 0.35.5) using a high-quality genome of the lysogenic *E. coli* K-12 MG1655 as reference (ERA18570755) (1). Next, we filtered the mutation predictions from breseq using the R statistical language (27) as follows: first, we removed mutations resulting from differences between our ancestor and the reference strain. Second, we filtered mutations so that only those present in populations that were treated with Amp, MMC or Amp + MMC remained. In addition, we assembled our reads into contigs using the de novo genome assembly algorithm of SPAdes (2). We used our genome assemblies with PHASTER (3, 4) to verify the presence/absence of phage Lambda+. Lastly, we used ResFinder (v 4.0) to search for antibiotic resistance genes in each sample (5, 6).

##### Whole Populations

Quality check and trimming was done as described for single clones. We examined the presence of mutations using breseq with the same reference as above (ERA18570755) using the polymorphism option. We exported breseq results into a table and extracted mutations related to phage resistance, chromosomal Amp-resistance (c_Amp^R^) and the frequency of Lambda+ carriage using the same method as for individual clones (Supp. Table 3).

### 4. Quantifying Candidate Mutations Over Time

Whole genome sequencing revealed three major types of genetic changes. This includes presence/absence of Lambda+ as well as mutations associated with phage and Amp resistance. Next, we analyzed the frequency and dynamics of these mutations over time. We used the same set of clones, i.e., 24 randomly selected clones from transfers 1-7, and 29 from two *s_lys* and *r_lys* populations that were either exposed to MMC or Amp+MMC, across analyses to generate a clear timeline of when phage loss as well as phage and antibiotic resistances emerged and spread in each population. We refer to this set of clones from here on as “candidate clones”.

#### Phage Loss

Whole genome sequencing revealed that clones that no longer produced free phage particles lost Lambda+. To infer the dynamics of this prophage loss we used clonal PCRs on our candidate clones with primers for Lambda+ as described above.

#### Phage Resistance

1. *Genomic Evidence:* First, we inferred the molecular mechanisms underlying phage resistance and their temporal dynamics. Whole genome sequencing revealed two different phage resistance mechanisms: (i) super-infection exclusion (SIE), where the presence of Lambda+ protects the host cell from additional infections, or (ii) surface-receptor mutations (SRM) where mutations in *MalT*, a protein involved in cell surface receptor production, prevent Lambda+ from binding to the host cell (28–32). To infer the nature of these two phage resistance mechanisms over time, we quantified the presence of Lambda+ and SRMs on our candidate clones.
2. *SRM*: Whole genome sequencing of single clones and whole populations revealed SRMs in *MalT*. Mutations in *MalT* come with a pleiotropic effect, i.e., a deficiency in the uptake of *malt*ose. This deficiency can be used to phenotypically infer phage resistance of individual clones using tetrazolium tetrachloride (TTZ) agar plates (29) on which we tested our candidate clones. As positive and negative controls, we used the Δ*MalT* strain from the KEIO collection and each respective ancestral clone (33).

We combined the two datasets, prophage and SRM presence, to construct the temporal dynamics of prophage loss and phage resistance and the molecular mechanism behind it, i.e., SIE or SRM.

#### Ampicillin Resistance

Whole genome sequencing revealed two trends regarding phage encoded ampicillin resistance (Amp^R^). (1) Under MMC selection, *r_lys* clones that lost their prophage also lost the phage encoded Amp^R^ and likely became susceptible to Amp (2) under Amp selection, populations previously susceptible to Amp acquired non-synonymous mutations in genes commonly associated with resistance to beta-lactam antibiotics (32–36). Therefore, we tested the phenotypic resistance to Amp of these two populations.

*r_lys populations*: To test whether *r_lys* populations that lost Lambda+ and its encoded Amp^R^, we performed a Minimum Inhibitory Concentration (MIC) assay using six clones from every evolved *r_lys* population subjected to all four selection regimes. We used Amp concentrations of 2, 4, 8, 16, 32, 64, 128, and 256 µg/ml to estimate the MIC and measured their OD600 at T0 and after 24 hours. We then used the difference between the two OD measurements to infer Amp resistance. A difference greater than 0.10 between the final and initial bacterial density indicated appreciable growth and thus resistance to Amp. To detect changes in the IC_50_, the concentration needed to inhibit 50% of the population, we fitted these growth data to a hill function (34,35).

*s_lys and n_lys populations*: To test, whether s_lys and n_lys populations subjected to Amp evolved chromosomal Amp-resistance (c_Amp^R^), we tested the MIC of six clones previously used for PCRs and TTZ assays in the same manner as the *r_lys* populations. To get a complete resistance profile of each population, we additionally tested the resistance of all 24 clones at the IC90 and IC99 concentrations of Amp. We grew each clone overnight at 37° C and re-inoculated them into 96-well plates containing one of two different concentrations of Amp corresponding to the IC90 and IC99 of the ancestral WT and measured OD600 at T0 and after 24 hours.

#### Ampicillin Removal

In the Amp+MMC treatment, we noticed a coexistence of phage carrying and phage free clones. This suggested that phages carrying the BLA-TEM resistance gene were cooperatively deactivating the Amp for phage free and thus antibiotic susceptible clones that did not have chromosomal mutations conferring Amp resistance. To confirm this, we tested the ability of lysogens carrying Amp^R^ to degrade Amp in liquid culture by performing an antibiotic removal test. To do so, we separately inoculated ancestral *n_lys*, *s_lys*, and *r_lys* bacterial populations containing the maximum concentration of Amp that *r_lys* bacteria can tolerate, i.e., 256 µg/ml and incubated them at 180 rpm overnight at 37° C. Cultures were then centrifuged at 5000 RPM for 10 minutes and the supernatant of each population was removed, and filter sterilized with a pore size of 0.2 mm. We plated and spotted the filtered supernatants to confirm that no viable bacteria respectively phages were present therein. Next, we inoculated the supernatant of the *r_lys* cultures, which we expected to contain beta-lactamase and the control supernatants of the *n_lys* and *s_lys* cultures with both the ancestral *n_lys* or ancestral *s_lys* populations, both of which are susceptible to Amp and incubated them overnight at 37° C at 180 rpm. We measured OD_600_ at inoculation and after 24 hours to measure bacterial growth in each of the supernatants.

#### Statistical analysis

All statistical analysis was done in R (v 4.1.3) using RStudio (v 2021.09.0). If not otherwise stated, we report mean(s) and standard error(s). For all analyses aimed to compare differences in free phages and bacteria we excluded control populations, i.e., *n_lys* populations as well as control treatments, i.e., LB and Amp. To compare temporal dynamics of phage and bacterial population sizes across the remaining genotypes and treatments we used a generalized least squares model (package *nlme* v3.1 – 160) with time, treatment, and genotype as well as all interactions as fixed effects. We used pairwise t-tests (package *stats*) to compare the change in phage population size from T8 to T29 in specific populations.

#### Stochastic model of prophage maintenance across environments

##### Model structure

We used a stochastic model of a well-mixed environment to simulate the evolutionary dynamics of Lambda+ lysogens, based on experimentally observed bacterial populations. We used one system of ordinary differential equations for each of the four treatment types with either type of bacterial lysogens, *N_s_lys_* and *N_r_lys_* (See Supplementary Information.docx). We do not explicitly model the free bacteriophage population because lysogen populations do not get (productively) superinfected. However, we approximate killing due to phages in MMC treatments for phage free and not phage resistant bacteria by a death rate β. The equations are based on population genetics models where populations grow at population-specific net growth rates *(g_x_)* and competition is implemented through a common carrying capacity (*K_total_)*. Bacterial cells also have a reduced growth rate *(g_s_lys,_ g_r_lys_)* when they are susceptible to phage attachment.

Depending on the treatment, bacteria die due to the killing effect of Amp (*ε)* or due to the killing caused by prophage induction through MMC *(α)*. The effect of a drug on bacterial growth, i.e., *ε* or *α,* is usually related to the concentration of a drug via a sigmoid curve (pharmacodynamic function). As the drug concentrations used in our model and LDA are in a regime far away from the MIC (minimal inhibitory concentration), we approximate the drug effect through a linear effect parameter. Subsequently, the concentration of Amp is given by *A* and the concentration of MMC by *M.* Both are decaying at rates δ*_A_* and δ*_M_* respectively, and Amp is degraded additionally in *r_lys* simulations with cooperative prophage-encoded resistance at a rate ψ and proportional to the population size of prophage carrying bacteria.

Here, we describe the genetic changes allowed in the most complex model variant, where antibiotic susceptible lysogens (*N_s_lys_*) are subjected to both MMC and Amp (see the Supplementary Information for details on adjustment of subpopulations and genetic changes in other model variants). We allow for three possible mutations to occur in *s_lys* populations in any order but without back-mutations: (i) chromosomal antibiotic resistance mutations at rate *μ_σ_* that confer complete resistance to the population (*N_σ_*), (ii) chromosomal surface receptor mutations conferring resistance to bacteriophage infection at rate μ*_λres_ (N_λres_* populations), which we assume increases the growth rate *(g_λres_)* of these cells as they are no longer negatively affected by repeated phage absorption (36), and (iii) loss of the prophage at rate μ_Δ*λ*_ (*N*_Δ*λ*_). Note, that prophage loss is usually caused by a large deletion, hence we assume this mutation rate to be lower than for phage and Amp resistance, which can be caused by one or few mutations.

For *r_lys* populations, we did not allow chromosomal resistance to arise as the lysogens were already Amp^R^. Emergence of chromosomal resistance was therefore only possible after a cell lost the prophage. We tested this assumption in an additional set of simulations, where we allowed chromosomal resistance to appear in any of the populations, but the pattern of prophage loss did not change substantially (Supp. Fig. 2).

**Figure 2:**
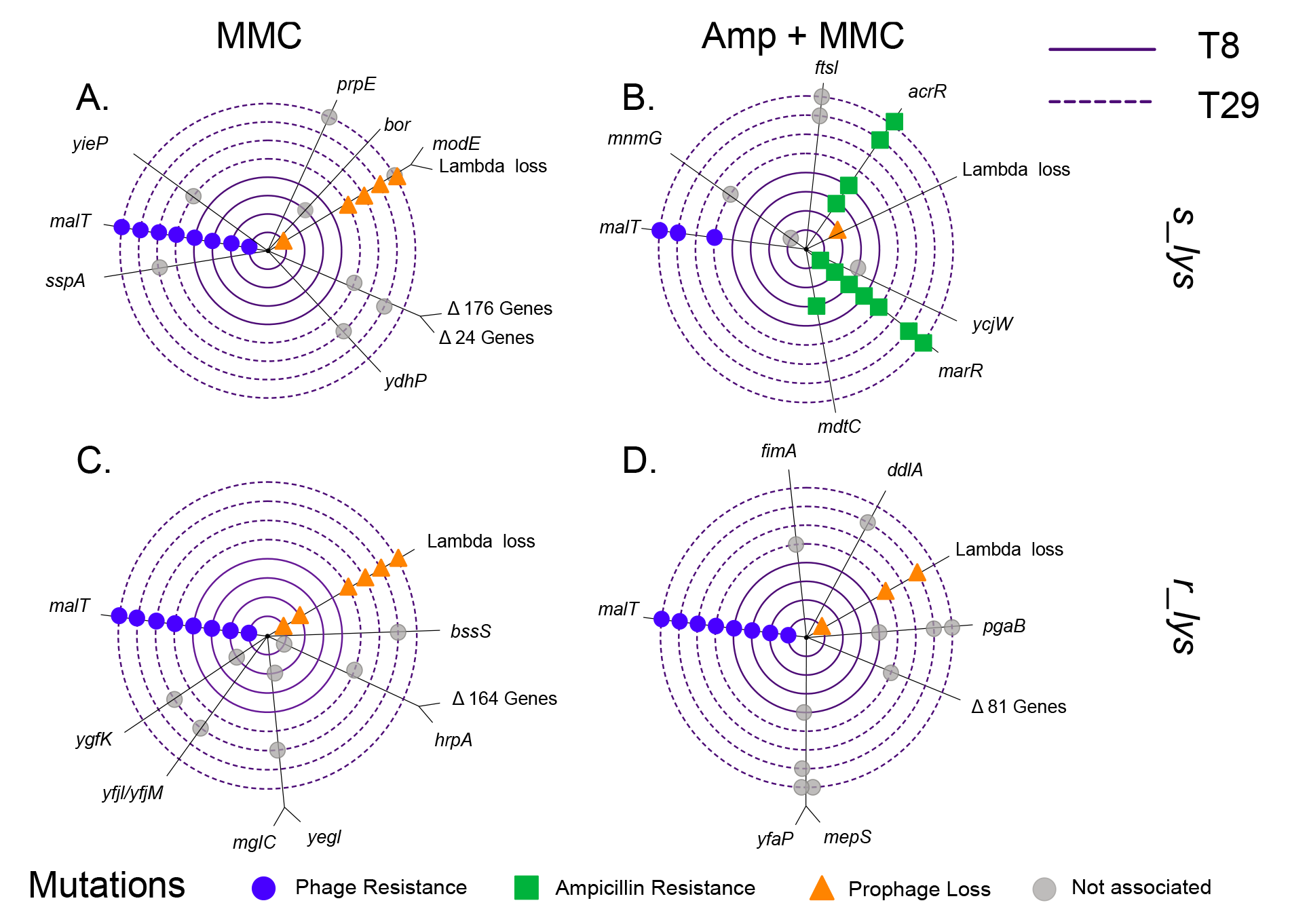
Genetic loci under positive selection as indicated by parallel genomic evolution in s_lys (top) and r_lys (bottom) populations exposed to MMC (left) or Amp+MMC (right). Each concentric circle corresponds to a single clone; Inner circles (solid) correspond to clones from T8, outer circles (dashed) correspond to clones from T29. Each colored point corresponds to one mutation event on the respective clone. Mutations associated with ampicillin resistance (*marR* and *acrR*) are shown in green, mutations associated with bacteriophage resistance (*MalT*) in blue, and deletion of Lambda+ in orange. Mutations that could not be associated with the different phenotypes are shown in grey. For more detailed information on the underlying mutations see Supp. Table 1. For more detailed information on population frequencies see Supp. Fig. 10.

In order to investigate the effect of cooperativity in the phage-encoded benefit, we performed model simulations where the prophages carried antibiotic resistance that provided purely an individual, non-cooperative benefit and compared it with the simulations where prophages provided cooperative antibiotic resistance. The non-cooperative benefit was achieved by removing the additional Amp degradation by resistant lysogens (i.e., setting γ to 0).

Details of equations and populations used in different model versions were adjusted as appropriate to the experimental setup and are detailed in Supplementary Information. We simulated the ODE model stochastically in R 3.6.0 using the *adaptivetau* package (27,37). We simulated this model for 50 transfers, and after each transfer 1% of the population was chosen randomly (R function *sample*) to make up the starting population of the next transfer. We started each transfer again with the chosen initial concentration of Amp and MMC. Based on empirical observations Amp and MMC affect bacterial population size only after a set time-period, which we included in our model as a delayed input time for Amp and MMC. We used the *tidyverse* suite of packages to take the averages of the 100 runs for each version of our model and plot them (38).

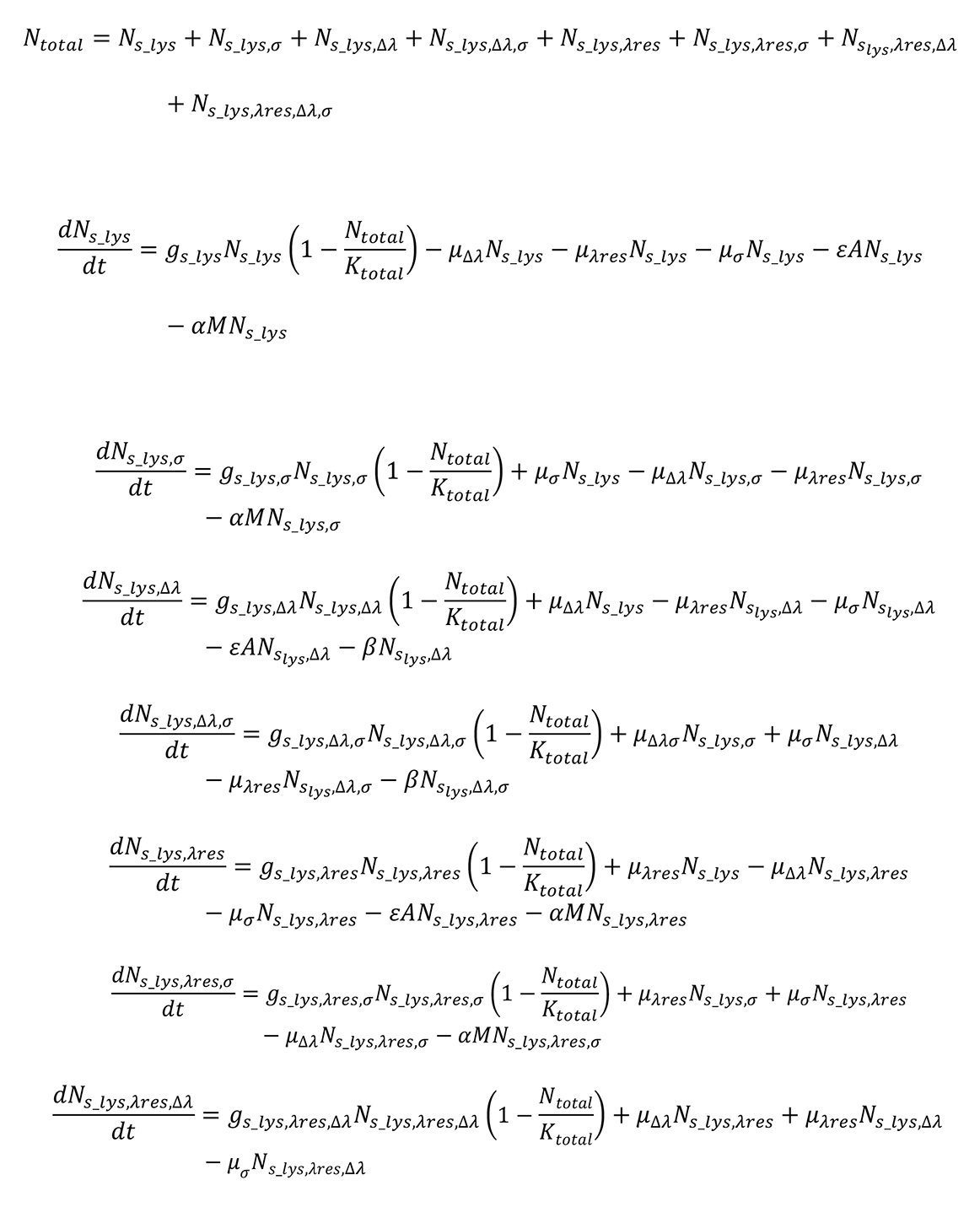

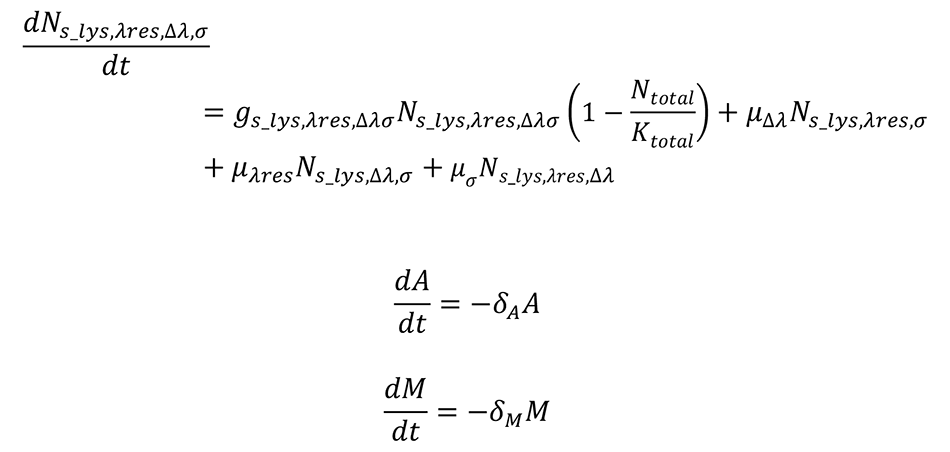

#### Model fitting

We fitted individual parameters of the model using a maximum-likelihood based method in R (*fminsearch* in the *Pracma* package) based on a deterministic version of our model. We fitted the growth rate of both *s_lys* and *r_lys* populations to a growth curve of the ancestral lysogen grown in nutrient rich media. We also fitted the effects of the two drugs Amp and MMC (*ε* and *α*) to growth curves of *s_lys* treated separately with MMC or Amp at the same concentration as in the selection experiment. To fit the mutation rates, we used data on the frequency of each type of mutation estimated from 24 clones per population per transfer for the first seven transfers of the experiment. These mutation rate approximations are in agreement with published mutation rates of *E. coli* genes (39). Using the fitted parameter set within our stochastic model gave a qualitatively good fit between the model and our empirical data (Supp. Fig. 3). For a list of parameter values used in the simulations and the parameter sensitivity analysis (LDA) see Supplementary Table 3 and Supplementary Table 4.

**Figure 3:**
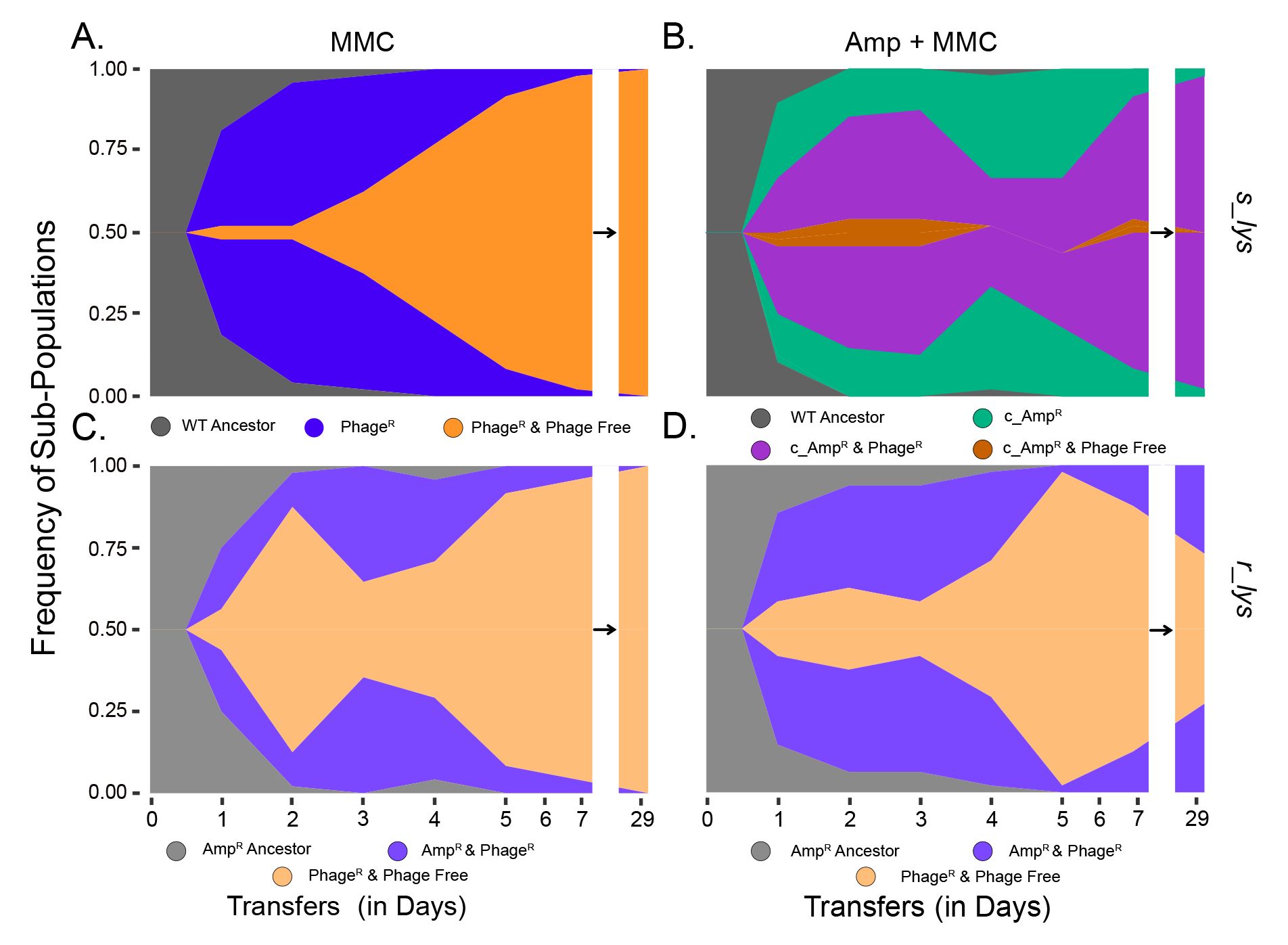
Muller plot showing sub-population frequencies over time of a single replicate population from the selection experiment. Colored areas are proportional to the frequencies of each sub-population in the population: ancestral WT (gray), phage resistant lysogens (blue), Amp^R^from chromosomal mutations (green), phage resistant and c_Amp^R^mutations (purple), phage resistant and phage free (orange/brown). Plots were generated by combining phenotypic and genotypic assays on sets of 24 individual clones per population for one replicate from each treatment and genotype combination.

#### Parameter sensitivity analysis (Linear Discriminant Analysis)

We tested the sensitivity of our simulation results to variations in the magnitude of drug effects and to variations in mutation rates using Latin hypercube sampling and Linear Discriminant Analysis (LDA) as we expect mutation rate and drug concentration to be influential in the loss of prophages (40). We created 10,000 sets of parameters using Latin hyper-cube sampling (R package *lhs* (41)) to evenly cover the sampling space for variation in five parameters: mutation rate to phage resistance *(μ_λres_)*, mutation rate to Amp-resistance (*μ_σ_*), mutation rate to prophage loss (*μ_Δλ_*), the effect of MMC (*α*), and the effect of Amp (*ε*). We used parameter values for mutations rates that evenly covered either side of our fitted values, but we used Amp and MMC effect values which were centered on a smaller value than our fitted one as higher values led to a high percentage of simulations where all populations die out. We ran 100 replicate simulations of each set of parameters for 50 transfers, at which point the system has reached steady state (as tested for a subset of parameters).

The results were categorized into different classes and analyzed using LDA in R (*lda* from the package *MASS* (42). The classes were based on prophage presence within the population, resulting in a majority phage containing class (> 51%), a majority phage free class (> 51%) and a mixed class where neither phage containing, nor phage free bacteria were dominant. We plotted an overview of the relative amounts of these classes as bar plots (Fig. 4A, Supp, Fig. 3) by utilizing a subset of 3335 parameter sets of the 10,000 parameter sets that had the same parameter values (2000 parameter sets for the simulations where chromosomal Amp-resistance mutations were allowed in *r_lys* bacteria). LDA was then used to determine which parameters separate these classes maximally. The magnitude and direction of the parameter arrows show their significance in separating certain classes, which correlates with the magnitude of their impact on the simulation results (Supp Fig. 8).

**Figure 4:**
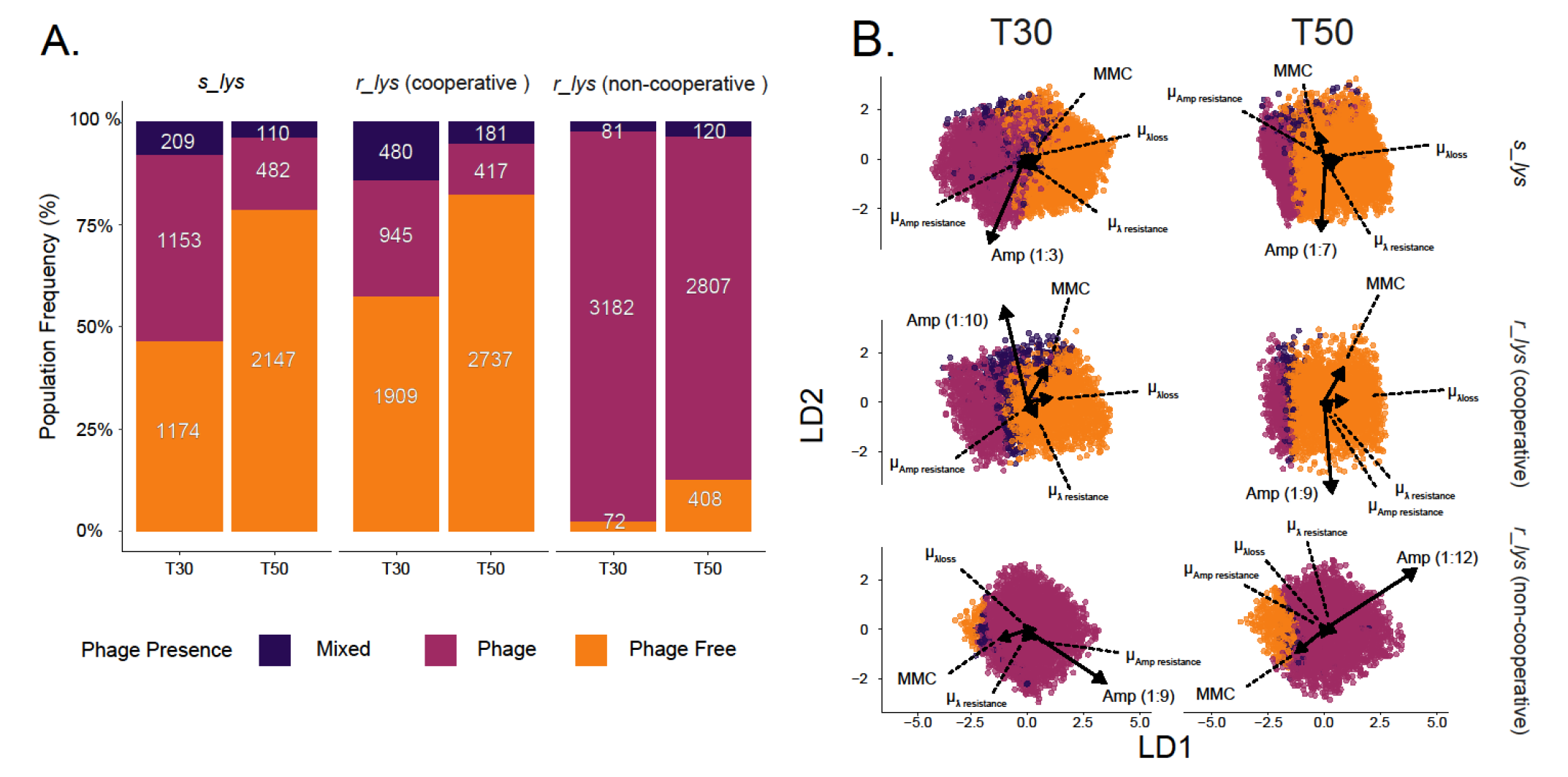
Frequency of bacteria from our simulations that maintain their prophage in the Amp+MMC treatment. Results from model simulations where we perform 100 simulation runs for each set of parameters with the Amp+MMC treatment for either *s_lys* or *r_lys* bacteria. We classified the simulation results based on whether the majority (>51%) of the population is phage free (orange), phage containing (amaranth purple), or the mixed (violet) with no majority. (A) Stacked bar plot of 3335 sets of parameter values that varied the mutation rates and effect of both Amp and MMC within the simulation, but where parameter values were the same across the different simulation scenarios. We compare these same parameter sets for *s_lys* bacteria, *r_lys* bacteria that carry cooperative antibiotic resistance, and *r_lys* bacteria that carry non-cooperative antibiotic resistance. (B) Linear Discriminant Analysis determining the impact of the three mutation rates as well as the two environmental effects: Amp, and MMC on the frequency of prophage loss. LD1 and LD2 are the principal axes of the LDA. Arrows show the magnitude and direction of a parameter in separating the classes. The effect size of Amp, for example, was generally much larger than all other factors (rescaling of Amp to fit the plot is shown in brackets).

The results of the LDA were used to visualize the relative impact of the two selection pressures MMC and Amp on prophage maintenance at T30 and T50 (Figure 4). For our visualizations, we removed simulations where every sub-population becomes zero. For the LDA plots, we averaged the frequency of prophage-carriers over 100 different simulations using the same parameter set of mutation rates, MMC and Amp effect values. We also visualized our results with contour plots that show the frequency of prophage carrying populations at a particular combination of Amp and MMC effect. For our contour plots, we averaged the frequency of prophage carriers over all simulations done for a parameter combination of MMC and Amp values (meaning that mutation rate parameters were not necessarily the same between simulations used for averaging).

## Results

To study the evolutionary dynamics of prophage maintenance in lysogen populations, we evolved three distinct *E. coli* genotypes: *n_lys* (non-lysogens), *s_lys* (lysogens carrying a Lambda+ prophage), and *r_lys* (lysogens carrying a Lambda+ prophage that encodes a beta-lactamase gene providing ampicillin resistance (Amp^R^)) in four different selection regimes for 30 serial transfers. These regimes imposed varying selection pressures on prophage carriage: (1) a control regime (nutrient rich broth), (2) a conflicting-selection regime that selected against the prophage (broth with mitomycin C (MMC) to increases prophage induction), (3) a selection regime for the prophage-encoded Amp^R^ (broth with Amp), and (4) a selection regime against the prophage but for its encoded Amp-resistance gene (broth with both Amp and MMC).

### Production of free phages varies across lysogen genotypes and selection regimes

First, we compared the ecological dynamics of bacteria and free phages between genotypes and treatments. The overall number of bacteria was reduced on transfer 1 (T1) in populations exposed to Amp and/or MMC. However, population sizes recovered after the first week and stabilized at levels comparable to the control (Supp. Fig. 4, 5) suggesting rapid adaptation to the different selection regimes.

In contrast, phage densities varied significantly among selection regimes (significant interaction in gls-model for treatment, genotype, and time: F_1,328_ = 26.55, p < 0.0001; Supp. Fig. 6, Supp. Fig. 7). In the absence of MMC, all lysogen populations produced a maximum phage titer of 10_5_ PFUs/ml on T1 before declining to 0 PFUs/ml (Supp. Fig. 6). This dynamic is not surprising because introduction into nutrient rich media can initially lead to an increase in spontaneous induction before the number of free phages stabilized at approximately ∼10_-8_ inductions per generation (43,44).

In the absence of Amp but the presence of MMC, phage production increased from an average of 1.4×10_6_ PFUs/ml at T1 to a maximum of 9.1×10_7_ PFUs/ml on T8 (Fig 1) before significantly declining (significant Time effect in gls-model: F_1,52_ = 4.97, p = 0.03, Fig 1), irrespective of phage genotype (non-significant genotype effect in gls-model: F_1,52_=0.64, p = 0.43), over the remaining experiment by an average of 3.5 orders of magnitude. In extreme cases, i.e., in two *r_lys* populations from T22 and one *s_lys* population from T15 onwards, we observed zero phage production.

This overall reduction in phage production seen under MMC selection contrasts sharply with populations exposed to the competing selective pressures, Amp+MMC. While the number of free phages in all *r_lys* populations remained stable over time (no significant time effect in gls-model: F_1,28_ = 0.443, p > 0.05; Fig1D), we observed strong fluctuations and no clear trend, across *s_lys* populations (Fig 1B).

At the end of the experiment, we tested the control populations exposed to LB, for the presence of prophages using an MMC induction assay. The assay confirmed that all *n_lys* populations were indeed phage free, whereas *r_lys* and *s_lys* populations remained inducible by MMC at the end of the experiment.

### Decline in phage particles at the population level is driven by individual cells

The overall decline in free phages under MMC selection could be caused by a reduction in average phage production across the entire population, or by individual cells within a given population that did not produce any phage particles. To differentiate between these two possibilities, we quantified phage particle production of 24 randomly selected clones from two replicate populations subjected to either MMC or MMC + Amp from timepoints with high (T8) or low (T29) levels of free phages. Overall, phage particle production followed a bimodal distribution (Supp. Fig. 8B) with one local maximum at zero, indicating no phage particle production, and a second local maximum at 1.64×10^6^ ± 2.93×10^5^ PFUs/ml, which is close to the average level from the start of the experiment at 9.14×10^5^ ± 3.19×10^5^ PFUs/ml (Supp. Fig. 8B).

In the MMC selection regime, we observed, irrespective of the underlying genotype, two different dynamics of free phage presence: in one population, the population size of free phages decreased from its maximum on T8 to zero by T29 (Supp. Fig. 8). In contrast, the other tested population did not produce any phages at both time points. These observations are in stark contrast to the Amp+MMC selection regime, where phage production remained stable within a given population over time (*s_lys* t = 5.825, p = 1 and *r_lys* t = 0.30123, p = 0.618; Supp. Fig. 8).

This bimodal distribution of phage particle production across all populations suggests that the decline in free phages under MMC selection can be explained by individual cells that have lost the ability to produce free phages.

### Prophage loss coincides with phage and antibiotic resistance mutations

To elucidate the molecular mechanisms underlying the absence of phage particle production, we sequenced two clones from two *s_lys* and two *r_lys* populations exposed to MMC or Amp+MMC, and two clones from one *s_lys* and one *r_lys* control population at T8 and T29. From these, we identified three main sets of genetic changes, each comprising one or more genetic loci, which can account for the observed phenotypic changes and ecological dynamics in the selection experiment. Strong parallel signatures observed across populations for each of these three groups indicate adaptive evolution to the respective selection regime. To determine the final frequencies of each mutation within every population, we additionally sequenced whole populations from the last transfer of the experiment.

The first set of genetic changes involves molecular resistance of bacterial cells to Lambda+ adsorption through surface receptor mutations (SRM; Figure 2). This set includes *malT,* which is involved in the uptake and utilization of maltose (45), and confers resistance to Lambda+ (28,30). In lysogens exposed to MMC and Amp+MMC, the frequency of non-synonymous single nucleotide variants (SNVs), insertions, and non-sense mutations within *malT* increased from 75% of the sampled clones on T8 to 100% on T29 (Fig 2 A). To test whether these mutations confer phage resistance through SRMs, i.e., maltose receptor inactivation, we assessed the ability of 24 individual clones from two replicate populations from T1-T5, T7 and T29 in *maltose* uptake by using a TTZ assay (29). This assay revealed that the frequency of SRMs rapidly increased to above 75% within the first 48 hours and reached a frequency of 100% in all *s_lys* and *r_lys* populations treated with MMC and Amp+MMC by T29 (Supp. Fig. 9).

The second genetic change involves variations in the presence or absence of Lambda+ (Fig. 2). In the absence of Amp but presence of MMC, Lambda+ was lost from all sequenced clones. This was supported by our population level sequencing data, which suggested that Lambda+ was present in less than 1% of the lysogenic populations from this treatment (Fig 2, Supp. Fig. 10). This stands in sharp contrast to the Amp+MMC selection regime, where Lambda+ persisted at nearly 100% in every *s_lys* population or ranged from ∼23% to 100% across *r_lys* populations (Supp. Fig. 10).

The third set of genetic changes includes various genes linked to Amp-resistance and appeared solely in the Amp+MMC selection regime. This set includes single nucleotide variants, deletions and insertions in four different loci: the multidrug resistance genes *marR* and *mdtH*, the multidrug-efflux pump *acrAB-tolC*, and the outer membrane gene *ompF* (52–55 Figure 2, Supp Table 2). Mutations in *marR* or *acrAB-TolC* reached fixation in three out of four *s_lys* populations (Figure 2A, Supp Table 2). An antibiotic susceptibility (MIC) assay confirmed that these mutations, which are commonly associated with resistance to beta-lactam antibiotics (32–36), confer resistance to the ampicillin concentration used in the selection experiment (Supp. Fig. 11) and explain the significant increase in final population sizes of initially Amp susceptible populations in the selection experiment over time (Effect of Amp on population sizes in gls: F = 120.6; p < 0.01, Supp. Fig. 4,5).

Although not visible at the clonal level our population level sequencing revealed that also two out of six *r_lys* populations exposed to Amp+MMC acquired mutations associated with Amp-resistance (Supp. Fig. 10). These two populations exhibited the highest frequency in prophage loss among all six-replicate *r_lys* populations, i.e., over 75% of the clones lost Lambda+ and its encoded Amp^R^. This suggests that the chromosomal Amp^R^ (c_Amp^R^) mutations compensate for the loss of the phage encoded Amp^R^ in *r_lys* populations subjected to the Amp+MMC selection regime.

Taken together, the genetic changes observed in this study indicate strong parallel adaptation of *s_lys* and *r_lys* populations to frequent prophage induction - via phage resistance mutations and prophage loss - and Amp selection – via chromosomal Amp-resistance mutations.

### Frequent prophage induction selects for rapid prophage loss characterized by distinct selective sweeps

As the loss of prophages occurred together with phage and Amp-resistance mutations, we next inferred the temporal dynamics of these genetic changes. We found that both *r_lys* and *s_lys* populations adapted to MMC through two selective sweeps: During the first sweep, lysogens acquired SRMs that prevented reinfections by free phages. Once SRM lysogens reached a frequency of ∼75%, this genotype was replaced by fixation of a phage-free SRM (Figure 3 A+C), resulting in the second selective sweep. This suggests strong selection for SRM when lysogens are exposed to frequent prophage induction.

To better understand the selective processes underlying prophage loss under MMC selection, we implemented our empirical observations into a stochastic model of lysogen evolution. Our simulations showed that the increased induction rate under MMC selection, coupled with a growth cost of phage adsorption to lysogens, are sufficient to produce the same two consecutive selective sweeps observed in the experiment: emergence of phage resistance followed by prophage loss (Supp. Fig. 3).

### Prophage encoded benefits and conflicting selective forces prolong prophage maintenance

In contrast to selection in the absence of Amp, *s_lys* populations subjected to Amp+MMC exhibited different evolutionary dynamics. Here, the first adaptive mutations observed were chromosomal Amp-resistance (c_Amp^R^; Fig. 3B), which suggests that in our experiment, the selection pressure imposed by Amp was stronger than the selection pressure imposed by MMC. Only after the emergence of c_Amp^R^ did these *s_lys* populations acquire SRM. Subsequently, this new genotype, which is resistant to Amp and the burden of phage absorption, rapidly spread through the population, reaching a frequency of nearly 100%. Surprisingly, although phage-free c_Amp^R^ variants did emerge, they did not increase in frequency as observed under MMC selection.

The need for three consecutive selective sweeps instead of two in *s_lys* populations under Amp+MMC selection suggests that prophage loss in these populations may require a longer timescale. To confirm this hypothesis, we simulated the evolution of *s_lys* populations under Amp+MMC selection over a wider timeframe. While our simulations revealed a mean frequency of prophage carriage of 99% on T30, prophage carriage decreased to 65% on T50. This demonstrates that, despite being costly, prophages are maintained longer in the presence of a second selective pressure because two consecutive selective sweeps - c_Amp^R^ and phage resistance - must occur before prophage-free cells can emerge (Figure 4A; 4B).

In contrast, *r_lys* populations exposed to Amp+MMC acquired c_Amp^R^ less frequently and instead underwent the same two selective sweeps as in the absence of Amp: emergence of phage resistance followed by phage loss. However, in the presence of both selection pressures, phage-free SRMs did not outcompete SRM *r_lys* lysogens, but instead co-existed at varying frequencies over time (Fig. 3D). This co-existence is likely explained by the presence of cooperative resistance. Specifically, the beta lactamase enzyme secreted by prophage-carrying *r_lys* cells can break down the Amp in the culture, allowing phage-free cheaters to arise and take advantage of their enzyme-producing kin. We confirmed the ability of *r_lys* bacteria to protect phage-free bacteria from an otherwise lethal concentration of Amp using an ampicillin removal assay (Supp. Fig. 12). This suggests that the selective advantage of the prophage-encoded cooperative Amp-resistance allows the prophage to persist in environments that would otherwise select for prophage loss.

Our simulations confirmed that the presence of cooperative resistance encoded on the prophage in our *r_lys* populations weakens the selection pressure for c_Amp^R^. As a result, prophages are more quickly lost in *r_lys* compared to *s_lys* populations. When prophages encode a non-cooperative benefit such as a drug-efflux pump, our simulations predict an even longer maintenance of prophages in lysogen populations (Figure 4A). The non-cooperative benefit prevents phage-free cheaters from emerging as they will not be protected from Amp killing by the other lysogens. This suggests, that in environments that otherwise select against prophages, conflicting selection hinders prophage loss as chromosomal resistance is required for competitive growth in these conditions - similar to what we observed in *s_lys* populations.

### Environmental factors trump mutation rates in determining prophage maintenance

The evolutionary dynamics of prophage loss showed environment-dependent distinct successions of selective sweeps. To determine the extent to which these successions depend on the availability of each genetic change, i.e., the variation in mutation rates of phage and antibiotic resistance and prophage loss, and the effect sizes of individual selection pressures, i.e., MMC and Amp, we used a Linear Discriminant Analysis (LDA). Our results revealed that mutation rates for phage and antibiotic resistance, and prophage loss had little impact on the outcome of our simulations. Instead, irrespective of the lysogen genotype, Amp had by far the strongest effect on the evolutionary trajectories, followed by MMC (Figure 4B).

For *s_lys* populations, prophage loss frequencies varied widely across the entire MMC-Amp parameter space, reflecting the differences in phage production between replicate populations in the selection experiment (Fig. 1B, Supp. Fig. 13). Our LDA explains this variability by demonstrating that Amp had the greatest impact on prophage maintenance despite not selecting for or against the prophage in this scenario (Figure 4B). That is because Amp substantially decreased the population size, reducing mutational input, and because selection for c_Amp^R^ was stronger than selection against the prophage. However, once c_Amp^R^ became more prevalent, population sizes increased again, and the selection pressure regarding prophage maintenance exerted by Amp became neutral.

In contrast, for *r_lys* bacteria, the frequency of prophage loss was primarily determined by the relative effect size of selection pressure for the phage-encoded benefit (Amp) and against the phage itself (MMC), as well as the duration of selection (Figure 4, Supp. Fig. 13). *r_lys* populations with a cooperative resistance gene lost the phage more quickly and to a greater extent than *s_lys* populations and this effect increased with increasing selection against prophages (Figure 4A, Supp. Fig. 13). However, when the resistance gene on the phage was non-cooperative, prophage loss in *r_lys* populations was significantly lower than in *s_lys* populations (Figure 4A). Specifically, selection by Amp outweighs MMC selection for non-cooperative *r_lys* populations across a range of sub-MIC values for both Amp and MMC, whereas MMC is the greater selection pressure for phage loss in cooperative *r_lys* bacteria (Supp. Fig. 13).

In summary, our findings suggest that prophage maintenance is influenced more strongly by complex selection patters resulting from environmental factors rather than mutational availability.

## Discussion

Active prophages are key players in the ecology and evolution of bacterial populations, with important consequences for higher order ecological interactions (50). While the benefits of prophages to their hosts are well established, they also pose a constant risk of death to their host bacterium. Here, we investigated the role of environmental conditions in determining the maintenance and loss of prophages, considering the fitness benefits and costs associated with active prophages.

We found that prophage maintenance and loss is primarily determined by environmental conditions that alter the fitness benefits or costs of an active prophage. In situations where the environment solely selects against the prophage, lysogenic populations can lose the prophage rapidly. However, prophage-encoded benefits and the presence of multiple selection pressures maintain prophages even when they entail substantial fitness costs. For instance, prophage-encoded resistance genes, which offer non-cooperative benefits can prolong prophage maintenance, as prophage loss in the absence of chromosomal drug resistance reduces bacterial fitness. In this sense, prophage loss can be thought of as crossing a fitness valley and the rise of chromosomal mutations that confer drug resistance may be slow since they are either selectively neutral or costly (51–53) if the same benefit is already provided by the prophage. However, when the benefits are cooperative, prophages are maintained at low frequencies because their gene product can protect phage-free cheaters that are likely to emerge if selection against prophages is high. Furthermore, the presence of an additional, unrelated selective pressure can maintain prophages by reducing population size and selecting more strongly for other beneficial mutations.

Our genomic analysis revealed that prophage loss occurs through environment-specific successions of selective sweeps. During the first sweep, lysogenic bacteria acquired surface receptor mutations (SRM) providing protection against phage infections. This enabled the second sweep, during which phage-free SRM rapidly increased in frequency. The rapid spread of SRM in lysogenic populations may seem surprising, as the integrated phage itself already protects the bacterium from superinfection. However, it can be explained by a repeated physiological burden of phage adsorption to the cell surface (36). Preventing phage adsorption via SRM might thus be a beneficial strategy in the presence of high phage titers (54,55) and allows for subsequent prophage loss, which ameliorates the costs of frequent induction.

Our simulations demonstrated that prophage maintenance and loss trajectories are primarily driven by environmental selection pressures, rather than mutational availability. This suggests that prophage loss is not simply a matter of chance mutations occurring in the bacterial genome. Rather, it is influenced by a complex interplay between environmental factors and specific genetic changes. By highlighting the strength of environmental factors relative to mutation rates in determining prophage loss trajectories, we suggest that bacterial populations may be able to adapt to changing environments in complex ways that are not necessarily predictable based solely on genetic changes.

The maintenance of active prophages is potentially analogous to the plasmid paradox (56–58). Plasmids should be lost from a population due to their high maintenance costs but are frequently found in bacterial populations because they can carry beneficial genes, such as resistances to antibiotics or heavy metals (56–58). Like plasmids, the net fitness effect of prophages is highly environment dependent but their ability to kill and lyse the host bacterium imposes an additional conflict for their hosts. Despite this death risk, which can be very costly for a lysogen population, we show that active prophages can be maintained within bacterial populations if they carry beneficial genes that confer a fitness advantage to their host. This can be thought of as a cooperation between the prophage and the bacterium, which may be further strengthened by the ability to deploy free phages as weapons during bacterial warfare (5,59).

Our study highlights the complex interplay between genetic and environmental factors that shape the dynamics of prophage maintenance and loss. We demonstrate the intricate selection pressures that act on mobile genetic elements, including the importance of accessory genes encoded thereon. Our results emphasize the need to consider the sociality of accessory genes in understanding the selective forces that act on prophages and their hosts. Given the omnipresence of prophages in the bacterial world, our findings have important implications for our understanding of bacterial evolution in changing environments. By gaining a deeper understanding of the interplay between these factors, we can improve our ability to predict and manage the evolution of microbial populations and the genetic elements within them.

