## Supplementary Figures for "Prophage maintenance is determined by environment-dependent selective sweeps rather than mutational availability"

**Supplementary Figure 1**: 20-hour growth curves of a lysogenic population in the presence of different concentrations of MMC ranging from 0µg/ml (top left) to 1.0µg/ml (bottom right). The concentration of MMC used in the selection experiment is underlined, i.e., 0.1 µg/ml.

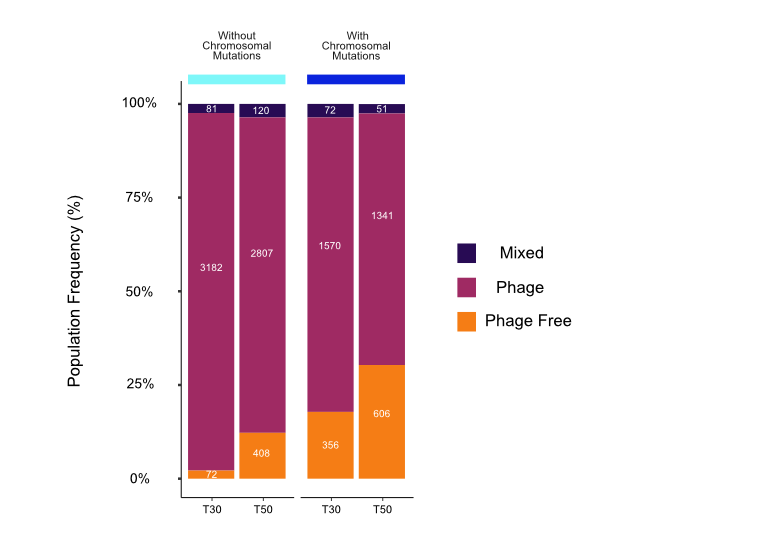

**Supplementary Figure 2:** Stacked bar plot showing the frequency of phage free or phage containing bacteria in the simulation data of non-cooperative *r_lys* bacteria. Frequencies are obtained as averages of 100 simulation runs per parameter set of the Amp+MMC treatment with r_lys bacteria for either 3335 (teal lines) or 2000 (blue line) parameter sets (varying mutation rates and effect of both Amp and MMC, see LDA) that had the same parameter values across model variations (Fig. 4A). In the simulations, chromosomal resistance mutations were not allowed (teal lines) or allowed (blue line) to occur in already resistant lysogens. We classified populations into ‘phage free’ or ‘phage’ if more than 51% of the population did not or did carry phages, respectively. Populations with no majority were considered mixed.

**Supplementary Figure 3:** Muller plots obtained from model simulations based on a parameter set fitted from empirical data and literature. Here our simulations are run for the equivalent of 50 transfers compared to the 29 transfers in our empirical study. A complete table of parameter values is given in Supplementary Table 3. The top row shows *s_lys* simulated populations and the bottom row *r_lys* simulations. The left column is the treatment regime of MMC alone and the right column is Amp and MMC together. The initial wild type for both genotypes is gray, and the ancestral bacteria is taken over by phage resistant (Phage^R^) bacteria (blue/purple) or chromosomally ampicillin resistant (c_Amp^R^) bacteria (green). Phage resistant bacteria is then displaced by phage free and phage resistant bacteria (orange), and in bacteria treated with Amp+MMC phage resistant, phage free and c_Amp^R^ bacteria appear (dark orange).

**Supplementary Figure 4:** 24-growth curves of each replicate population using sub-samples of the plates used in the main experiment every seven transfers for each replicate population of genotype and treatment from the selection experiment. The general trend is towards higher OD values over time especially after the first seven transfers in populations treated with Amp and/or MMC.

**Supplementary Figure 5**: Bacterial dynamics, i.e., bacterial cell counts derived from flow cytometry over the course of the selection experiment. Each line represents a replicate population within that combination of genotype (rows) and treatment (columns).

**Supplementary Figure 6:** Free phage particles, i.e., plaque forming units (PFUs/ml) measured once a week during the selection experiment. This shows the number of phages per replicate population present in the culture of each genotype (rows) by treatment (columns) combination.

**Supplementary Figure 7:** Phage particle production shown as the number of bacteriophages (PFUs, Supp. Fig. 6) relative to bacterial cell counts (Supp. Fig. 5) for each genotype (rows) by treatment (columns) combination.

**Supplementary Figure 8**: (A) The number of free phages that 24 randomly selected clones from two populations of both MMC and Amp + MMC treated populations of both s_lys and r_lys genotypes produced when induced with 0.1 µg/ml of MMC. We took 24 clones from T8 and T29 to observe changes in the ability to produce free phages across time between individual clones. (B) Stacked histograms of the same data from T8 and T29 respectively to demonstrate the distribution of free phage production. The red dotted line represents the average number of free phages from populations treated with MMC on the first transfer of the selection experiment.

**Supplementary Figure 9**: Indication of phenotypic resistance of evolved bacteria to bacteriophage lambda measured from TTZ (Tetrazolium-Tetra Chloride Agar) plates. This shows the frequency of maltose mutations for one population (Pop2) from each genotype (rows) and treatment (column) combination.

**Supplementary Figure 10:** Each concentric circle represents the circular chromosome of an evolved population. Each pie chart shows the frequencies of a specific mutation within a given population. Different shades of the same color represent different mutations at the same locus. Mutations that are not associated with Amp or phage resistance or the loss of prophages are removed for clarity. All mutations are listed in Supplementary Table 2.

**Supplementary Figure 11**: Reaction norms demonstrating the change in IC50 (in µg/ml) from the beginning (Transfer 0) to the end (Transfer 29) of the selection experiment. Each line corresponds to a replicate population that was tested for their Amp resistance; some lines overlap as they have the same IC_50_ values.

**Supplementary Figure 12:** The growth of (A) *n_lys* and (B) *s_lys* bacteria susceptible to ampicillin in four different types of supernatant originating from: no bacteria (red), *n_lys* bacteria (green), *s_lys* bacteria (purple), or *r_lys ­*bacteria (blue). The x-axis denotes the initial concentration of Amp that was added to the culture before the inoculation with bacteria and 24-hour incubation from which the supernatant was taken.

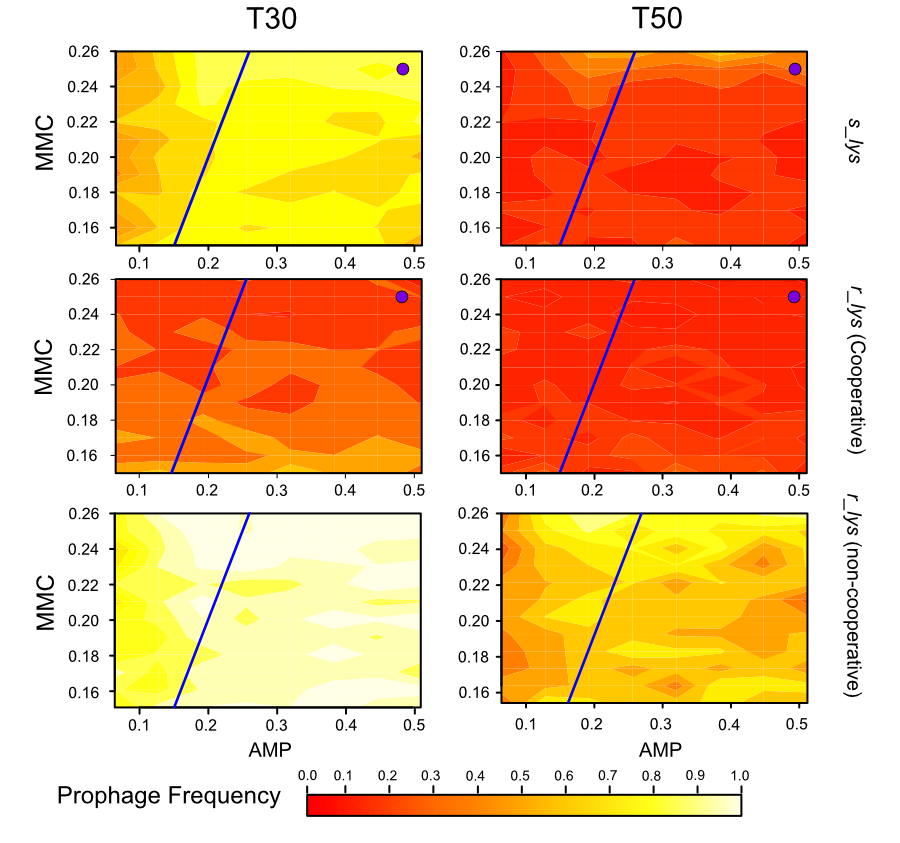

**Supplementary Figure 13:** Contour plot showing the average frequency of prophage carrying bacteria in simulated populations for different MMC and Amp effect values. The frequency was averaged over all the parameter sets that share the same Amp and MMC value (but they contain different mutation rates). Here we use 10000 parameter sets for the cooperative *r_lys* and *s_lys* simulations contour plots and 3335 parameter sets for non-cooperative *r_lys* simulations. We compare here both the difference between *s_lys* and *r_lys* simulated populations and *r_lys* bacteria with cooperative or non-cooperative antibiotic resistance. Here the effect of Amp and MMC pertains to the killing capacity of the drug (population killed/ unit time) times the concentration of that drug (µg/ml). The blue solid line indicates where the selective pressure of MMC is equal to that of Amp. The purple dots are the MMC and Amp equivalent of what is utilized in the selection experiment.

### **Supplementary Table 3**: Fitted Model Parameter values

| **Parameter** | **Description** | **Estimated Value** | **Source** |
| --- | --- | --- | --- |
| g*_slys_* , g_rlys_ | Growth rate of ancestral lysogens | 0.48 | Parameter Fitting |
| g_slys,λres_ | Growth rate of phage resistant lysogens | 0.52 | Parameter Fitting |
| g_slys,Δλ_ | Growth rate of phage free bacteria | 0.48 | Parameter Fitting |
| g_slys,λres,Δλ_ | Growth rate of phage resistant and phage free bacteria | 0.56 | Parameter Fitting |
| K_total_ | Carrying Capacity of Bacteria | 1.77×10^9^ | Parameter Fitting |
| E (ε) | Effectiveness of Ampicillin | 0.077 | Parameter Fitting |
| I (⍺) | Effectiveness of Mitomycin C | 2.53 | Parameter Fitting |
| δ_A_ | Degradation of Ampicillin | 0.034 | Parameter Fitting |
| δ_M_ | Degradation of Mitomycin C | 0.0065 | Parameter Fitting |
| τ_A_ | Input Time Ampicillin | 0.62 | Parameter Fitting |
| τ_M_ | Input Time Mitomycin C | 3.57 | Parameter Fitting |
| μ_λres_ | Mutation rate - Resistance to lambda phage | 1×10^-7^ | Literature (1,2), Parameter Fitting |
| μ $\sigma$ | Mutation rate - Resistance to Ampicillin | 1× 10^-6^ | Parameter Fitting |
| μ_Δλ_ | Mutation rate - Loss of Lambda Phage | 1×10^-9^ | Parameter Fitting |

1.	Drake, J. W. A constant rate of spontaneous mutation in DNA-based microbes. *Proc. Natl. Acad. Sci.* **88,** 7160–7164 (1991).

2.	Drake, J. W., Charlesworth, B., Charlesworth, D. & Crow, J. F. Rates of spontaneous mutation. *Genetics* **148,** 1667–1686 (1998).

### **Supplementary Table 4**: Range of parameter values used in our set of 10,000 parameters.

| **Parameter** | **Description** | **Starting Value** | **Ending Value** |
| --- | --- | --- | --- |
| E (ε) | Effectiveness of Ampicillin | 0.01 | 0.08 |
| I (⍺) | Effectiveness of Mitomycin C | 1.5 | 2.6 |
| μ_λres_ | Mutation rate - Resistance to lambda phage | 1×10^-9^ | 1×10^-5^ |
| μ $\sigma$ | Mutation rate - Resistance to Ampicillin | 1×10^-9^ | 1×10^-5^ |
| μ_Δλ_ | Mutation rate - Loss of Lambda Phage | 1×10^-11^ | 1×10^-8^ |

### **Mathematical model variants (List of Equations)**

Here we describe all variants of the differential equations used in the mathematical model simulations. Parameter values used are presented in Supplementary Tables 3 and 4.

The sum of all sub populations (if present in a given model variant)

$$N_{total}=N_{slys}+N_{slys,\lambda res}+N_{slys,\Delta\lambda}+N_{slys,\sigma}+N_{slys,\lambda res,\sigma}+N_{slys,\lambda res,\Delta\lambda}+N_{slys,\sigma,\Delta\lambda}+N_{slys,\lambda res,\sigma,\Delta\lambda}$$

**Control (in LB Broth):**

The population of ancestral “normal” lysogens

$$\frac{{dN}_{slys}}{dt}=g_{slys}N_{slys}\left( 1-\frac{N_{total}}{K_{total}} \right)-\mu_{\Delta\lambda}N_{slys}-\mu_{\lambda res}N_{slys}$$

Population cured for the lambda prophage

$$\frac{{dN}_{slys,\Delta\lambda}}{dt}=g_{slys,\Delta\lambda}N_{slys,\Delta\lambda}\left( 1-\frac{N_{total}}{K_{total}} \right)+\mu_{\Delta\lambda}N_{slys}-\mu_{\lambda res}N_{slys,\Delta\lambda}$$

Population resistant to bacteriophage Lambda

$$\frac{{dN}_{slys,\lambda res}}{dt}=g_{slys,\lambda res}N_{slys,\lambda res}\left( 1-\frac{N_{total}}{K_{total}} \right)+\mu_{\lambda res}N_{slys}-\mu_{\Delta\lambda}N_{slys,\lambda res}$$

Population resistant to bacteriophage lambda and cured of lambda prophage

$$\frac{{dN}_{slys,\lambda res,\Delta\lambda}}{dt}=g_{slys,\lambda res,\Delta\lambda}N_{slys,\lambda res,\Delta\lambda}\left( 1-\frac{N_{total}}{K_{total}} \right)+\mu_{\Delta\lambda}N_{slys,\lambda res}+\mu_{\lambda res}N_{slys,\Delta\lambda}$$

***s_lys* in MMC:**

When subjected only to mitomycin C the differential equations for each population are given by:

The population of ancestral “normal” lysogens

$$\frac{{dN}_{slys}}{dt}=g_{slys}N_{slys}\left( 1-\frac{N_{total}}{K_{total}} \right)-\mu_{\Delta\lambda}N_{slys}-\mu_{\lambda res}N_{slys}-\alpha MN_{slys}$$

Population cured for the lambda prophage

$$\frac{{dN}_{slys, \Delta\lambda}}{dt}=g_{slys,\Delta\lambda}N_{slys,\Delta\lambda}\left( 1-\frac{N_{total}}{K_{total}} \right)+\mu_{\Delta\lambda}N_{slys}-\mu_{\lambda res}N_{slys,\Delta\lambda}-\beta N_{\Delta\lambda}$$

Population resistant to bacteriophage Lambda

$$\frac{{dN}_{slys,\lambda res}}{dt}=g_{slys,\lambda res}N_{slys,\lambda res}\left( 1-\frac{N_{total}}{K_{total}} \right)+\mu_{\lambda res}N_{slys}-\mu_{\Delta\lambda}N_{slys,\lambda res}-\alpha MN_{\lambda res}$$

Population resistant to bacteriophage lambda and cured of lambda prophages

$$\frac{{dN}_{slys,\lambda res,\Delta\lambda}}{dt}=g_{slys,\lambda res,\Delta\lambda}N_{slys,\lambda res,\Delta\lambda}\left( 1-\frac{N_{total}}{K_{total}} \right)+\mu_{\Delta\lambda}N_{slys,\lambda res}+\mu_{\lambda res}N_{slys,\Delta\lambda}$$

Degradation of Mitomycin C

$$\frac{dM}{dt}=-\delta_{M}M$$

***s_lys* in Amp:**

When subjected to ampicillin the differential equations for each population are given by:

The population of ancestral “normal” lysogens

$$\frac{{dN}_{slys}}{dt}=g_{slys}N_{slys}\left( 1-\frac{N_{total}}{K_{total}} \right)-\mu_{\Delta\lambda}N_{slys}-\mu_{\lambda res}N_{slys}-\mu_{\sigma}N_{slys}-\varepsilon AN_{slys}$$

The population of “normal” lysogens with chromosomal ampicillin resistance ($\sigma$)

$$\frac{{dN}_{slys,\sigma}}{dt}=g_{slys,\sigma}N_{slys,\sigma}\left( 1-\frac{N_{total}}{K_{total}} \right)+\mu_{\sigma}N_{slys}-\mu_{\Delta\lambda}N_{slys}-\mu_{\lambda res}N_{slys}$$

Population cured for the lambda prophage

$$\frac{{dN}_{slys,\Delta\lambda}}{dt}=g_{slys,\Delta\lambda}N_{slys,\Delta\lambda}\left( 1-\frac{N_{total}}{K_{total}} \right)+\mu_{\Delta\lambda}N_{slys}-\mu_{\lambda res}N_{slys,\Delta\lambda}-\mu_{\sigma}N_{slys,\Delta\lambda}-\varepsilon AN_{slys,\Delta\lambda}-\beta N_{slys,\Delta\lambda}$$

Population cured for lambda prophage and resistant to Ampicillin

$$\frac{{dN}_{slys,\Delta\lambda,\sigma}}{dt}=g_{slys,\Delta\lambda\sigma}N_{slys,\Delta\lambda,\sigma}\left( 1-\frac{N_{total}}{K_{total}} \right)+\mu_{\Delta\lambda}N_{slys,\sigma}+\mu_{\sigma}N_{slys,\Delta\lambda}-\mu_{\lambda res}N_{slys,\Delta\lambda,\sigma}-\beta N_{slys,\Delta\lambda,\sigma}$$

Population resistant to bacteriophage Lambda

$$\frac{{dN}_{slys,\lambda res}}{dt}=g_{slys,\lambda res}N_{slys,\lambda res}\left( 1-\frac{N_{total}}{K_{total}} \right)+\mu_{\lambda res}N_{slys}-\mu_{\Delta\lambda}N_{slys,\lambda res}-\mu_{\sigma}N_{slys,\lambda res}-\varepsilon AN_{slys,\lambda res}$$

Population resistant to bacteriophage Lambda and ampicillin

$$\frac{{dN}_{slys,\lambda res,\sigma}}{dt}=g_{\lambda res,\sigma}N_{slys,\lambda res,\sigma}\left( 1-\frac{N_{total}}{K_{total}} \right)+\mu_{\lambda res}N_{slys,\sigma}+\mu_{\sigma}N_{slys,\lambda res}-\mu_{\Delta\lambda}N_{slys,\lambda res,\sigma}$$

Population resistant to bacteriophage lambda and cured of lambda prophage

$$\frac{{dN}_{slys,\lambda res,\Delta\lambda}}{dt}=g_{\lambda res,\Delta\lambda}N_{slys,\lambda res,\Delta\lambda}\left( 1-\frac{N_{total}}{K_{total}} \right)+\mu_{\Delta\lambda}N_{slys,\lambda res}+\mu_{\lambda res}N_{slys,\Delta\lambda}-\mu_{\sigma}N_{slys,\lambda res,\Delta\lambda}-\varepsilon AN_{slys,\lambda res,\Delta\lambda}$$

Population resistant to bacteriophage lambda, cured of lambda prophage and resistant to ampicillin

$$\frac{{dN}_{slys,\lambda res,\Delta\lambda,\sigma}}{dt}=g_{\lambda res,\Delta\lambda,\sigma}N_{\lambda res,\Delta\lambda,\sigma}\left( 1-\frac{N_{total}}{K_{total}} \right)+\mu_{\Delta\lambda}N_{\lambda res,\sigma}+\mu_{\lambda res}N_{\Delta\lambda,\sigma}+\mu_{\sigma}N_{\lambda res,\Delta\lambda}$$

Ampicillin degradation

$$\frac{dA}{dt}=-\delta_{A}A$$

***s_lys* in Amp and MMC:**

When subjected to mitomycin C and ampicillin the differential equations are given by:

The population of ancestral “normal” lysogens

$$\frac{{dN}_{slys}}{dt}=g_{slys}N_{slys}\left( 1-\frac{N_{total}}{K_{total}} \right)-\mu_{\Delta\lambda}N_{slys}-\mu_{\lambda res}N_{slys}-\mu_{\sigma}N_{slys}-\varepsilon AN_{slys}-\alpha MN_{slys}$$

The population of “normal” lysogens with resistance to ampicillin

$$\frac{{dN}_{slys,\sigma}}{dt}=g_{slys,\sigma}N_{slys,\sigma}\left( 1-\frac{N_{total}}{K_{total}} \right)+\mu_{\sigma}N_{slys}-\mu_{\Delta\lambda}N_{slys,\sigma}-\mu_{\lambda res}N_{slys,\sigma}-\alpha MN_{slys,\sigma}$$

Population cured for the lambda prophage

$$\frac{{dN}_{slys,\Delta\lambda}}{dt}=g_{slys,\Delta\lambda}N_{slys,\Delta\lambda}\left( 1-\frac{N_{total}}{K_{total}} \right)+\mu_{\Delta\lambda}N_{slys}-\mu_{\lambda res}N_{slys,\Delta\lambda}-\mu_{\sigma}N_{slys,\Delta\lambda}-\varepsilon AN_{slys,\Delta\lambda}-\beta N_{slys,\Delta\lambda}$$

Population cured for lambda prophage and resistant to Ampicillin

$$\frac{{dN}_{slys,\Delta\lambda,\sigma}}{dt}=g_{slys,\Delta\lambda,\sigma}N_{slys,\Delta\lambda,\sigma}\left( 1-\frac{N_{total}}{K_{total}} \right)+\mu_{\Delta\lambda}N_{slys,\sigma}+\mu_{\sigma}N_{slys,\Delta\lambda}-\mu_{\lambda res}N_{slys,\Delta\lambda,\sigma}-\beta N_{slys,\Delta\lambda,\sigma}$$

Population resistant to bacteriophage Lambda

$$\frac{{dN}_{slys,\lambda res}}{dt}=g_{slys,\lambda res}N_{slys,\lambda res}\left( 1-\frac{N_{total}}{K_{total}} \right)+\mu_{slys,\lambda res}N_{slys}-\mu_{\Delta\lambda}N_{slys,\lambda res}-\mu_{\sigma}N_{slys,\lambda res}-\varepsilon AN_{slys,\lambda res}-\alpha MN_{slys,\lambda res}$$

Population resistant to bacteriophage Lambda and ampicillin

$$\frac{{dN}_{slys,\lambda res,\sigma}}{dt}=g_{slys,\lambda res,\sigma}N_{slys,\lambda res,\sigma}\left( 1-\frac{N_{total}}{K_{total}} \right)+\mu_{\lambda res}N_{slys,\sigma}+\mu_{\sigma}N_{slys,\lambda res,\sigma}-\mu_{\Delta\lambda}N_{slys,\lambda res,\sigma}-\alpha MN_{slys,\lambda res,\sigma}$$

Population resistant to bacteriophage lambda and cured of lambda prophage

$$\frac{{dN}_{slys,\lambda res,\Delta\lambda}}{dt}=g_{slys,\lambda res,\Delta\lambda}N_{slys,\lambda res,\Delta\lambda}\left( 1-\frac{N_{total}}{K_{total}} \right)+\mu_{\Delta\lambda}N_{slys,\lambda res}+\mu_{\lambda res}N_{slys,\Delta\lambda}-\mu_{\sigma}N_{slys,\lambda res,\Delta\lambda}-\varepsilon AN_{slys,\lambda res,\Delta\lambda}$$

Population resistant to bacteriophage lambda, cured of lambda prophage and resistant to ampicillin

$$\frac{{dN}_{slys,\lambda res,\Delta\lambda,\sigma}}{dt}=g_{slys,\lambda res,\Delta\lambda,\sigma}N_{slys,\lambda res,\Delta\lambda,\sigma}\left( 1-\frac{N_{total}}{K_{total}} \right)+\mu_{\Delta\lambda}N_{slys,\lambda res,\sigma}+\mu_{\lambda res}N_{slys,\Delta\lambda,\sigma}+\mu_{\sigma}N_{slys,\lambda res,\Delta\lambda}$$

Ampicillin degradation

$$\frac{dA}{dt}=-\delta_{A}A$$

Degradation of Mitomycin C

$$\frac{dM}{dt}=-\delta_{M}M$$

***r_lys* in MMC**

Here the loss of prophage also means the loss of ampicillin resistance (𝜎).

When subjected only to mitomycin C the differential equations for each population appear as follows:

The population of ancestral “normal” lysogens

$$\frac{{dN}_{rlys}}{dt}=g_{rlys}N_{rlys}\left( 1-\frac{N_{total}}{K_{total}} \right)-\mu_{\Delta\lambda}N_{rlys}-\mu_{\lambda res}N_{rlys}-\alpha MN_{rlys}$$

Population cured for the lambda prophage

$$\frac{{dN}_{rlys,\Delta\lambda}}{dt}=g_{rlys,\Delta\lambda}N_{rlys,\Delta\lambda}\left( 1-\frac{N_{total}}{K_{total}} \right)+\mu_{\Delta\lambda}N_{rlys}-\mu_{\lambda res}N_{rlys,\Delta\lambda}-\beta N_{rlys,\Delta\lambda}$$

Population resistant to bacteriophage Lambda

$$\frac{{dN}_{rlys,\lambda res,\sigma}}{dt}=g_{rlys,\lambda res}N_{rlys,\lambda res}\left( 1-\frac{N_{total}}{K_{total}} \right)+\mu_{\lambda res}N_{rlys}-\mu_{\Delta\lambda}N_{rlys,\lambda res}-\alpha MN_{rlys,\lambda res}$$

Population resistant to bacteriophage lambda and cured of lambda prophage

$$\frac{{dN}_{rlys,\lambda res,\Delta\lambda}}{dt}=g_{rlys,\lambda res,\Delta\lambda}N_{rlys,\lambda res,\Delta\lambda}\left( 1-\frac{N_{total}}{K_{total}} \right)+\mu_{\Delta\lambda}N_{rlys,\lambda res}+\mu_{\lambda res}N_{rlys,\Delta\lambda}$$

Degradation of Mitomycin C

$$\frac{dM}{dt}=-\delta_{M}M$$

***r_lys* in Mitomycin C and Ampicillin:**

The loss of Lambda prophage also leads to the loss of ampicillin resistance (𝜎).

When subjected to mitomycin C and ampicillin the differential equations are given as follows:

The population of ancestral ampicillin resistant lysogens with resistance to ampicillin

$$\frac{{dN}_{rlys}}{dt}=g_{rlys}N_{rlys}\left( 1-\frac{N_{total}}{K_{total}} \right)-\mu_{\Delta\lambda}N_{rlys}-\mu_{\lambda res}N_{rlys}-\alpha MN_{rlys}$$

Population cured for the lambda prophage

$$\frac{{dN}_{rlys,\Delta\lambda}}{dt}=g_{rlys,\Delta\lambda}N_{rlys,\Delta\lambda}\left( 1-\frac{N_{total}}{K_{total}} \right)+\mu_{\Delta\lambda}N_{rlys}-\mu_{\lambda res}N_{rlys,\Delta\lambda}-\mu_{\sigma}N_{rlys,\Delta\lambda}-\beta N_{rlys,\Delta\lambda}-\varepsilon AN_{rlys,\Delta\lambda}$$

Population cured for lambda prophage and resistant to Ampicillin

$$\frac{{dN}_{rlys,\Delta\lambda,\sigma}}{dt}=g_{rlys,\Delta\lambda}N_{rlys,\Delta\lambda}\left( 1-\frac{N_{total}}{K_{total}} \right)+\mu_{\sigma}N_{rlys,\Delta\lambda}-\mu_{\lambda res}N_{rlys,\Delta\lambda,\sigma}-\beta N_{rlys,\Delta\lambda,\sigma}$$

Population resistant to bacteriophage Lambda with prophage-based resistance to ampicillin

$$\frac{{dN}_{rlys,\lambda res}}{dt}=g_{rlys,\lambda res}N_{rlys,\lambda res}\left( 1-\frac{N_{total}}{K_{total}} \right)+\mu_{\lambda res}N_{rlys}-\mu_{\Delta\lambda}N_{rlys,\lambda res}-\alpha MN_{rlys,\lambda res}$$

Population resistant to bacteriophage lambda and cured of lambda prophage

$$\frac{{dN}_{rlys,\lambda res,\Delta\lambda}}{dt}=g_{rlys,\lambda res,\Delta\lambda}N_{rlys,\lambda res,\Delta\lambda}\left( 1-\frac{N_{total}}{K_{total}} \right)+\mu_{\Delta\lambda}N_{rlys,\lambda res}+\mu_{\lambda res}N_{rlys,\Delta\lambda}-\varepsilon AN_{rlys,\lambda res,\Delta\lambda}-\mu_{\sigma}N_{rlys,\lambda res,\Delta\lambda}$$

Population resistant to bacteriophage lambda, cured of lambda prophage and resistant to ampicillin

$$\frac{{dN}_{rlys,\lambda res,\Delta\lambda,\sigma}}{dt}=g_{rlys,\lambda res,\Delta\lambda,\sigma}N_{rlys,\lambda res,\Delta\lambda,\sigma}\left( 1-\frac{N_{total}}{K_{total}} \right)+\mu_{\lambda res}N_{rlys,\Delta\lambda,\sigma}+\mu_{\sigma}N_{rlys,\lambda res,\Delta\lambda}$$

Ampicillin degradation (for communal (cooperative) resistance, ampicillin is actively degraded at a rate that is proportional to the resistant bacterial cells)

$$\frac{dA}{dt}=-\delta_{A}A-\gamma\left( N_{rlys}+N_{rlys,\lambda res} \right)A$$

Degradation of Mitomycin C

$$\frac{dM}{dt}=-\delta_{M}M$$
